# Topography, Spectral Characteristics, and Extra-to-Intracranial Propagation Pathways of EMG

**DOI:** 10.1101/808253

**Authors:** J. Lahr, L.D.J. Fiederer, O. Glanz, A. Schulze-Bonhage, T. Ball

## Abstract

**Objective:** Intracranial EEG (iEEG) plays an increasingly important role in neuroscientific research and can provide informative control signal for brain-machine interfaces (BMI). While it is clear that electromyographic (EMG) activity of extracranial origin reaches intracranial recordings, the topographic and spectral characteristics of intracranial EMG have been scarcely investigated. It is currently unclear how these characteristics compare to those of physiological brain activity. Little is also known about the exact pathways of extra- to intracranial volume conduction, including the role of craniotomy defects.

**Methods:** In 5 epilepsy patients under invasive pre-neurosurgical EEG monitoring, we examined chewing-related effects (ChREs) as a source of intracranial EMG activity and compared those effects with physiological brain activity of 9 patients during several behavioural tasks. These included speech production, finger movements, and music perception. Further, we analyzed the association of craniotomy defects (burr-holes and saw-lines) and the intracranial EMG-effects based on the individual post-operative images.

**Results:** ChRE presented with a spatially smooth distribution across almost all intracranial electrodes with the maximum below the temporal muscle. In contrast, the responses of neural origin were spatially more focalized. ChREs were broad-banded and had a higher spectral power and affected higher frequencies than event-related neural activity. ChRE were largely independent of the individual configuration of craniotomy defects. However, we found indications that the silicone sheet, in which electrocorticography (ECoG) electrodes are embedded, attenuates EMG influences, when sufficiently large.

**Conclusion:** The present work is the first comprehensive evaluation of topographic and spectral characteristics of EMG effects in iEEG based on a large sample of subjects. It shows that chewing-related EMG can affect iEEG recordings with higher power than typical physiological brain activity, especially in higher spectral frequencies. As the topographic pattern of ChRE is largely independent of the individual position of craniotomy defects, a direct pathway of volume conduction through the intact skull plays an important role for extra-to-intracranial signal propagation. Intracranial EMG activity related to natural behavior should be accounted for in neuroscientific and BMI applications, especially when based on high-frequency iEEG components. A detailed knowledge of EMG properties may help to design both EMG-reducing algorithms and ECoG grids with a high shielding factor.

**Highlights:**

- First comprehensive description of chewing-related EMG artifacts in iEEG recordings
- EMG artifacts and brain activity have distinct topographic and spectral iEEG characteristics
- Chewing EMG reaches the brain with higher spectral power than task-related brain activity
- Chewing-related EMG artifacts are largely independent of the the position of craniotomy defects

## 1. Introduction

Intracranially-measured electroencephalography (iEEG) is increasingly being used in neuroscientific research because of its superior signal quality over non-invasive electroencephalography (EEG; Ball et al., 2009c; Schalk, 2010). In addition, brain-machine interfaces (BMIs) for paralyzed patients have emerged as a new field for clinical application of iEEG (Ball et al., 2009c; Lahr et al., 2015; Leuthardt et al., 2006, 2004; Pistohl et al., 2008; Schalk et al., 2007; Wang et al., 2013; Xie et al., 2017). While artifact contamination has extensively been investigated in EEG (Dworetzky et al., 2010; Goncharova et al., 2003; Tong and Thankor, 2009), only few studies on artifacts in iEEG have appeared to date, such as those related to blinking (Ball et al., 2009a), saccades (Jerbi et al., 2009a; Kovach et al., 2011a), the heart cycle (Kern et al., 2013) and electromyographic (EMG) activity (Fiederer et al., 2016; Liu et al., 2004a; Otsubo et al., 2008a). Together, these previous studies demonstrate that such artifacts have an impact on iEEG recordings.

The previously-reported contamination of iEEG by artifacts, however, was often much more subtle than physiological neural effects observed using iEEG (e.g., Ball et al., 2009a), and they maximally reached the amplitude of actual neural responses (e.g., Kovach et al., 2011a). Hence, whether contaminating signals might become substantially larger than those of neural iEEG signals is still unclear.

Moreover, the exact conduction pathways by which this extra-to-intracranial signal propagation takes place have remained unknown. Due to the surgical procedure for the placement of iEEG electrodes, there are two different ways by which extracranial signals can enter the intracranial cavity. First, through craniotomy defects (burr holes, saw lines), and second, through the intact skull tissue. In the first case, intracranial artifacts would be expected to occur in a relatively focalized manner, underneath the craniotomy defects, constituting a ‘reverse breach effect’ (see the personal communication of J. Gotman cited in (Otsubo et al., 2008b)) that may be difficult to distinguish from the typically likewise focalized iEEG responses of neural origin. In the second scenario, intracranial contamination from extracranial sources would be expected to show a spatially widespread distribution, independent of the individual configuration of the craniotomy defects. This would be due to the spatial filtering properties of the skull (Neuling et al., 2012; Paul L Nunez and Srinivasan, 2006) – similarly to the well-known spatial blurring of EEG, which takes place in the opposite direction (Dannhauer et al., 2011; Lanfer et al., 2012b). In preliminary work we showed, using simulations in a single subject, that chewing-related artifacts likely propagate through the intact skull (Fiederer et al., 2016). Detailed empirical evidence on EMG pathwasy and the associated signal properties was, however, still lacking.

In the present study we address the following, hitherto unresolved questions: (i) How do intracranially recorded chewing-related effects (ChRE) effects differ from event-related brain activity with respect to topographic and spectral characteristics? (ii) Are the intracranial manifestations of EMG spatially focalized or diffuse? (iii) Does their overall topography depend on the individual configuration of the craniotomy defects resulting from electrodes implantation?

To address these questions, we investigated the intracranial manifestations of ChRE in 5 patients implanted with different iEEG configurations of electrodes and craniotomy defects. We compared these effects with sereral control conditions that presented physiological event-related brain responses during a variety of behavioral tasks (speech produciton, finger movements, and music perception) performed by 9 patients. We focused on ChREs as they constitute a potentially frequent artifact that may occur during natural behavior and that need to be accounted for in neuroscientific research and the development of clinical BMI applications. This is a particularly importand concern in the case of BMI applications for tetraplegic patients where self-feeding is a desired application (Collinger et al., 2013).

## 2. Methods

### 2.1 Intracranial EEG

#### 2.1.1 Subjects

Twelve subjects with medically intractable epilepsy who were under evaluation for neurosurgical treatment participated in the current study after having provided written informed consent. All subjects had subdurally implanted iEEG electrodes whose locations depended on the individual clinical requirements (see Table 1). For the ChRE analysis, we selected subjects who had at least one large ECoG grid with 8×8 contacts to be able to obtain extended and continuous topographies. iEEG electrodes were implanted for a period of 5 to 10 days. The ECoG electrodes were mounted on a flexible silicone substrate (Ad-Tech, Racine, WI) at a 10-mm center-to-center inter-electrode distance. They were made of stainless-steel or platinum discs which were 4 mm in diameter. In addition to ECoG grids and strips, penetrating depth electrodes in the hippocampus (1-mm diameter, 10 contacts with a 5-mm contact-to-contact distance) were also implanted in some subjects. The study was approved by the Ethics Committee of the University Hospital of Freiburg (Germany).

**Table 1:**
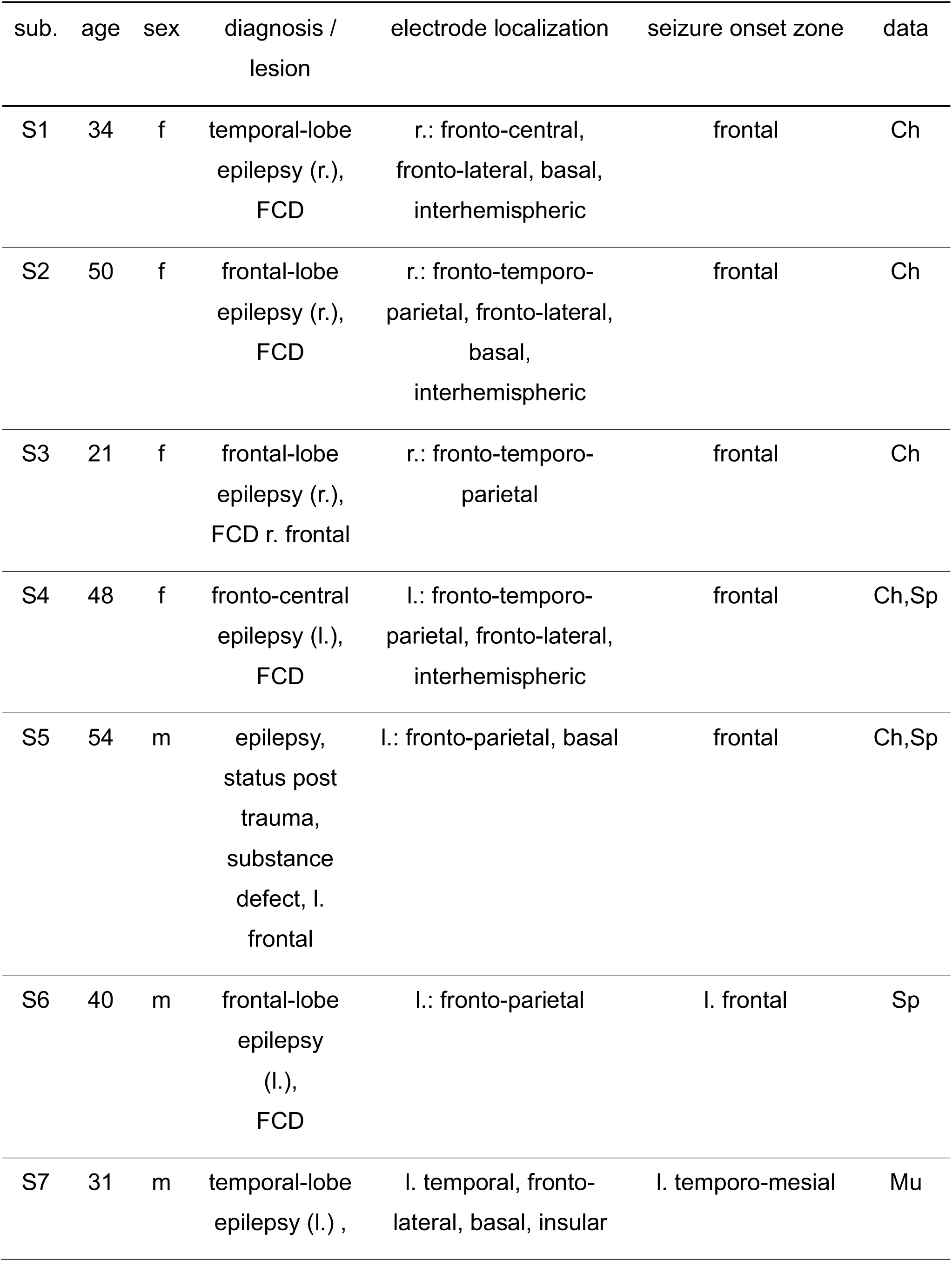
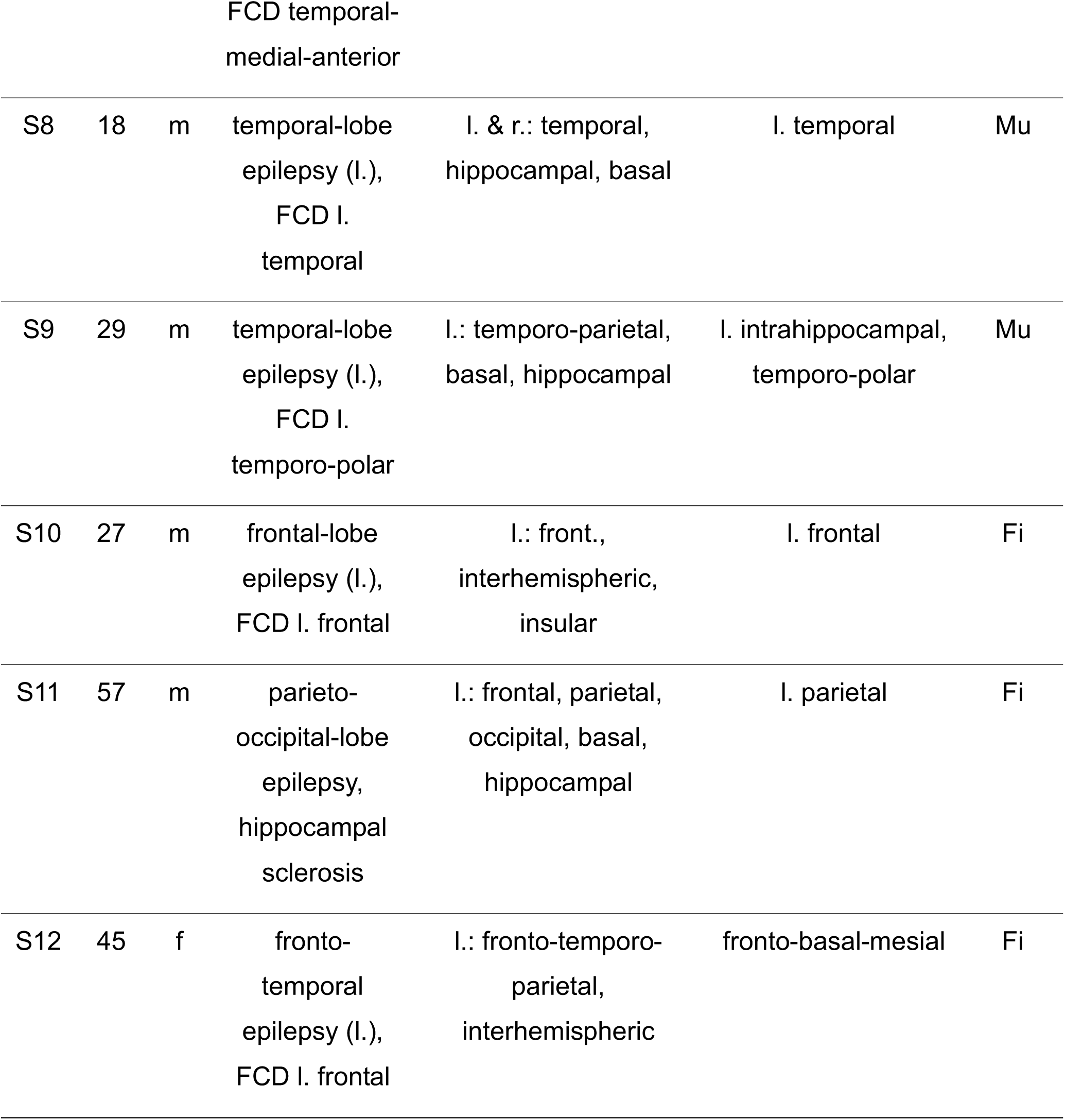
Subject overview. Abbreviations: sub.: subject; data.: analyzed datasets (Ch: chewing; Mu: Music perception; Fi: Finger movement; Sp: Speech production); m: male, f: female; FCD: focal cortical dysplasia; r.: right; l.: left

#### 2.1.2. Data acquisition

iEEG was recorded at a sampling rate of 1024 Hz using a clinical AC EEG-system (IT-Med, Usingen, Germany) in all subjects except for S12, in whom the sampling rate was 256 Hz. Digital video, synchronized with neural data, was recorded at 25 frames per second at VGA resolution. EEG according to the 10-20-system (Klem et al., 1999) was recorded simultaneously in all subjects in whom ChRE were analyzed. Electrodes with technical recording problems (e.g., due to broken wires) were excluded from all analyses.

### 2.2 Chewing-related Effects

#### 2.2.1 Trial selection for ChRE analysis

The onset and end of chewing-related EMG bursts were marked for each chewing event based on video- and EEG recordings (please refer to (Fiederer et al., 2016) for a detailed description of trial selection), and their arithmetic mean was defined as the 0-s time point for each trial. A total of 1652 trials was acquired (S1: 551 trials; S2: 438 trials; S3: 264 trials; S4: 252 trials; S5: 147 trials).

#### 2.2.2 Data analysis

EEG and iEEG data were separately re-referenced to a common average reference (CAR), as it is typically done in EEG and iEEG studies (Ball et al., 2009b; Canolty et al., 2007; Crone et al., 2006; Towle et al., 2007). Trials were cut from the continuous data from −2 s to 2 s with respect to the 0-s time point of the chewing event. Sliding-window fast fourier transformation was performed with a window length of 250 ms and a step width of 24.41 ms for the 1024-Hz data, and with a window length of 250 ms and a step width of 23.44 ms for the 256-Hz recordings. A baseline period (200 ms) centered between consecutive chewing events was defined to calculate the relative power. For the computation of relative power changes, the time-frequency spectra were divided by the median baseline power averaged across trials and then scaled logarithmically. Fig. 1 gives an overview of the ChRE time-domain and time-frequency domain data.

**Figure 1:**
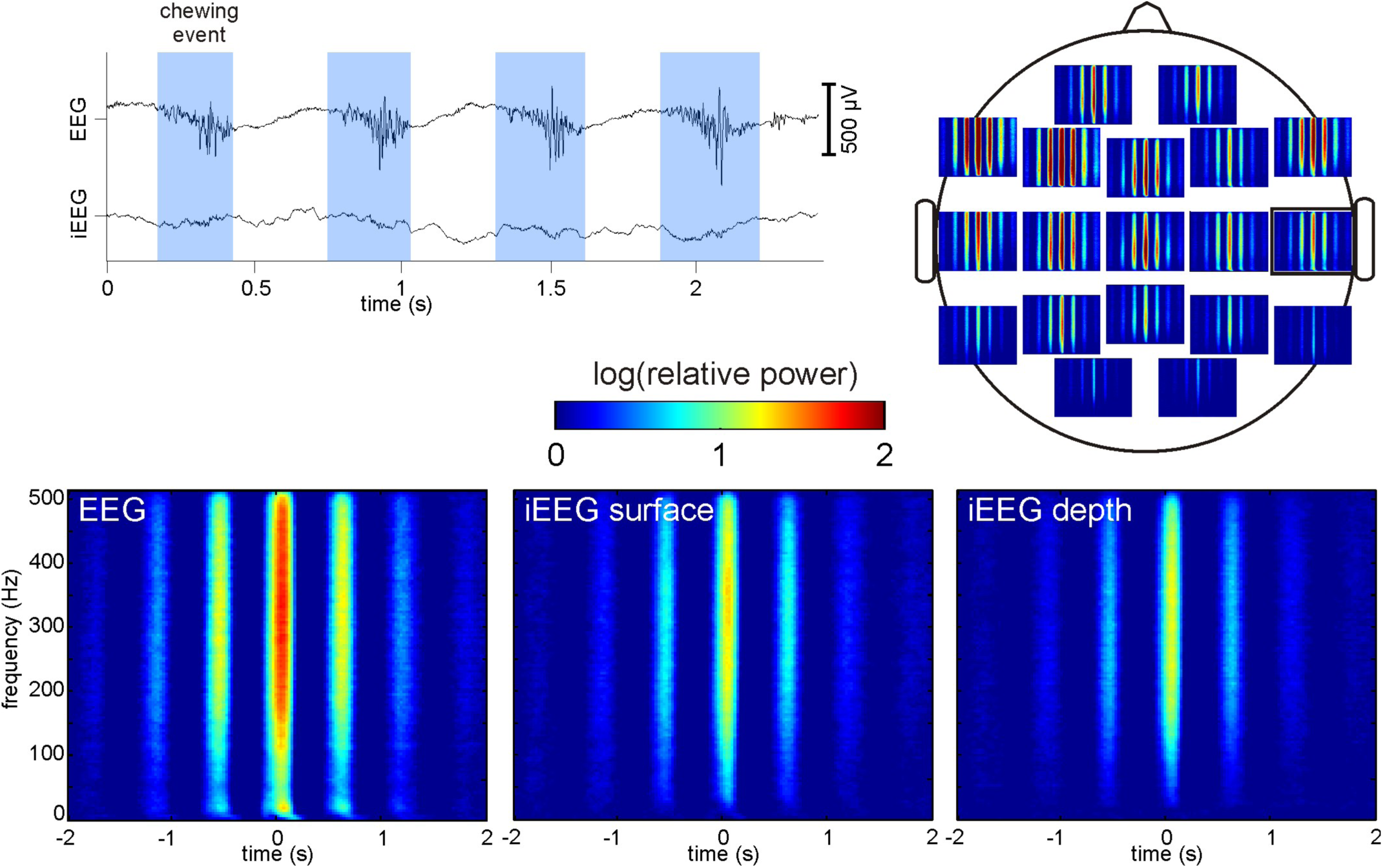
ChRE data overview. **(a)** Raw time-domain of concurrent EEG and iEEG data. ChREs are marked by the light-blue boxes. EMG activity can be clearly seen in the EEG trace. It is also visible in the iEEG trace albeit at a much lower amplitude. **(b)** ChRE time-frequency spectra topography in the EEG. Note the lower power of the ChRE over the right hemisphere, ipsilateral to the implantation cite. This pattern of weaker chewing ipsilateral of the implantation was seen across all 5 subjects. It is likely due to the incision made to the temporalis muscle and skin during surgery and the resulting pain during chewing. The contact marked by the black box is also shown in (c). **(c)** Comparison of the time-frequency spectra of EEG, surface iEEG (ECoG) and depth iEEG (stereo EEG).

#### 2.2.3 Relation of electrodes to craniotomy defects and anatomical landmarks

To determine the relation of the implanted electrodes to craniotomy defects (burr holes and saw lines) and to major anatomical landmarks (lateral and central sulci, further abbreviated as LS and CS, respectively), we used computed tomography (CT)-acquired images, X-ray data (Figs. 1 & 8), complemented by data from magnetic resonance imaging (MRI) (Figs. 3 & 4). These data were all acquired after electrode implantation in the individual subjects. As we focused on ChRE, the relation between the position of the temporal muscle and the recording sites was of particular interest. To relate the position of the temporal muscle to the position of the grid electrodes, the temporal line, the temporal muscle and the coronoid process of the mandible were reconstructed on the lateral x-ray images of the subjects. To this end, we took a vectorized image of the skull from an anatomy textbook (Gray, 1918) as a template. It was rotated and scaled in order to fit the individual anatomy of the subjects as represented by the lateral x-ray images. Then, craniotomy defects and electrode positions were determined and marked on the lateral x-ray image of the skull.

**Figure 2:**
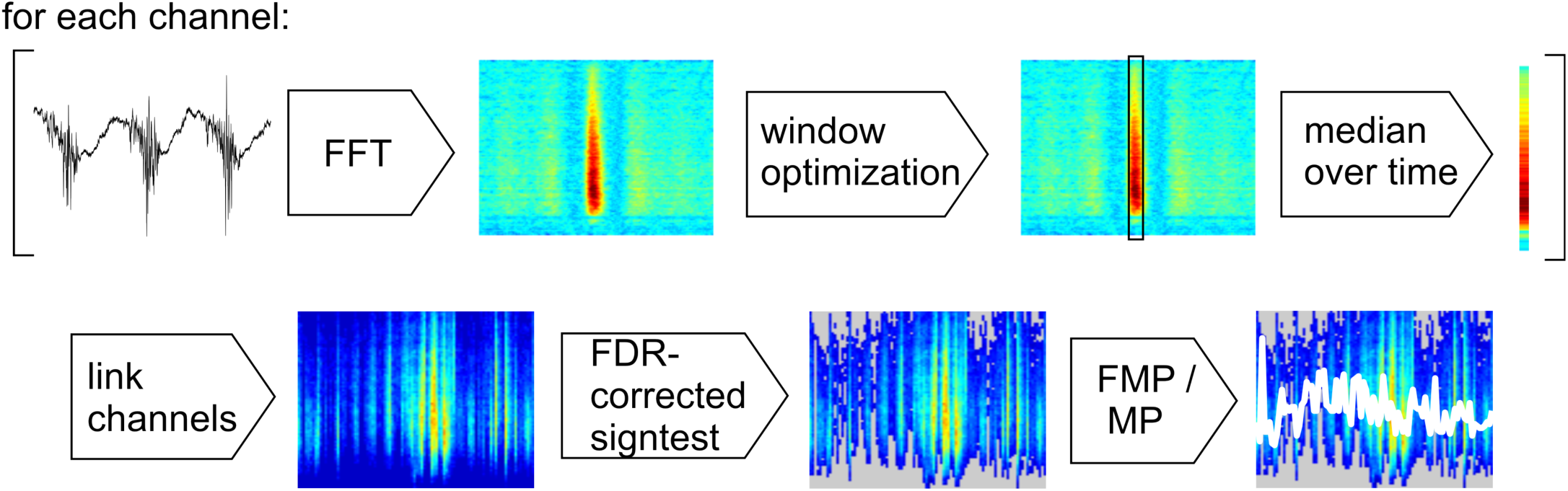
Flowchart of signal processing steps. Trials were processed using a sliding-window fast-Fourier transformation, resulting in a spectrogram. Then, the time-optimized windows were defined, and the median over time was calculated. The resulting frequency spectra of all channels were concatenated (visualized as vertical lines in the lower panels of this figure) and non-significant frequency bins were removed (visualized in grey). The result is an overview of the event-related frequency spectra of all electrodes in a subject. Finally, the MP and the FMP were extracted.

**Figure 3:**
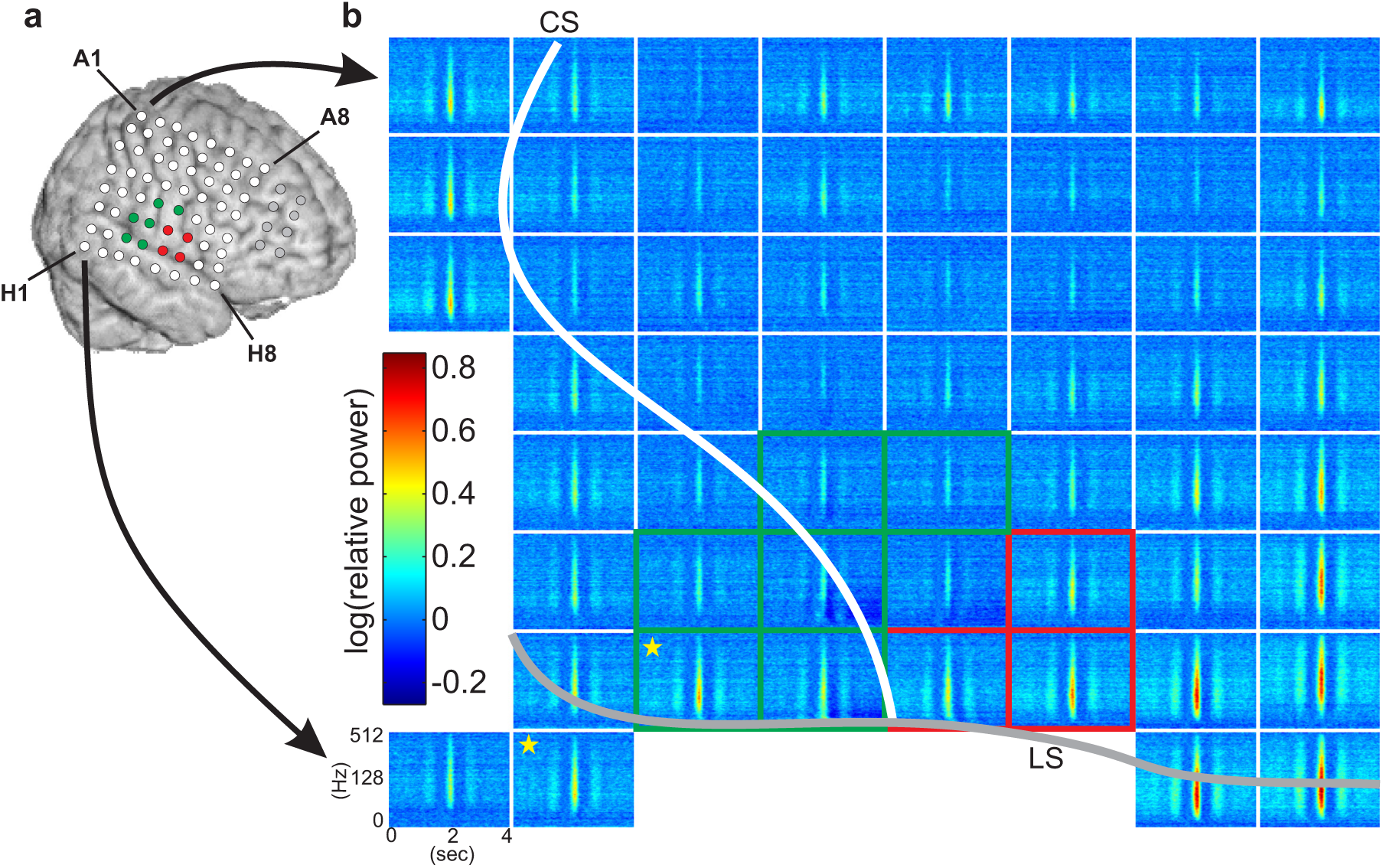
Intracranial topography of chewing-related ECoG power modulations. **(a)** 8×8 ECoG grid position in relation to the brain surface of S1, obtained from individual MRI data. During electrical cortical stimulation, electrodes in green and red displayed orofacial sensory and orofacial motor responses, respectively. A prefrontal 2×4 grid is shown in gray. The individual contacts of the electrode grid have labels (e.g., A1, H8) for ease of spatial reference. **(b)** Chewing-related spectrogram for all electrodes of the 8×8 grid (electrodes with technical recording problems were omitted and are therefore not visualized). Color encodes the logarithmic power change relative to the baseline. The orientation of the electrode grid is the same as in (a). Green and red boxes indicate electrodes with orofacial motor and sensory responses as in (a). The course of the lateral sulcus (LS) is depicted by a gray line and that of the central sulcus (CS) by a white line. Chewing-related power increases showed a spatially widespread distribution without any specific relation to the orofacial cortex. They crossed anatomical landmarks, e.g., bridging over the LS (indicated by yellow stars).

**Figure 4:**
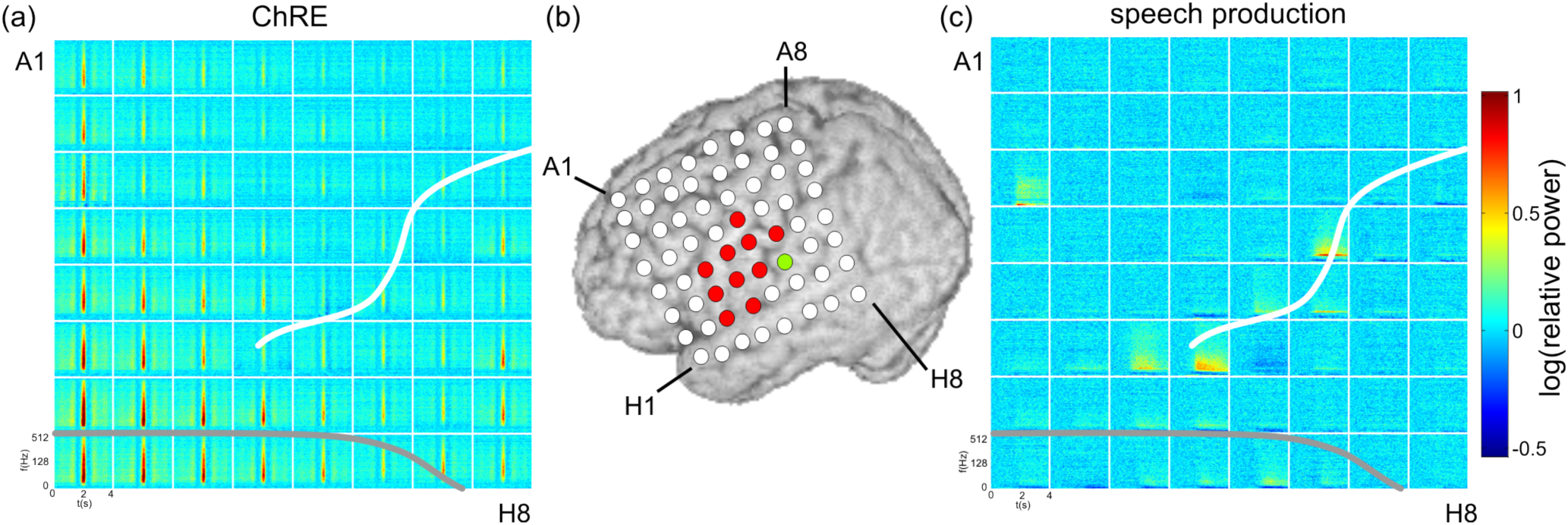
A comparison of the topographic distribution of chewing- and speech-related spectral responses (S4). **(a)** Spectrogram of ChREs. The course of the LS is depicted by a gray line, and the CS by a white line. Note the spatially widespread distribution bridging the LS. **(b)** ECoG grid position in relation to the brain surface obtained from individual subject’s MRI data. Electrodes with motor or sensory responses during the electrical cortical stimulation are marked in red and green, respectively. **(c)** Spectrograms obtained in the same subject during speech production showed spatially focalized responses in areas predominantly in or adjacent to the ESM-defined mouth sensorimotor cortex (orange outline). Conventions as in (a). In contrast to the ChREs in (a), speech-production-related responses were most prominent in frequencies below 200 Hz and showed an accompanying power decrease in lower frequencies (dark blue, approx. 8 to 32 Hz). (a) & (b) adapted from (Fiederer et al., 2016).

**Figure 5:**
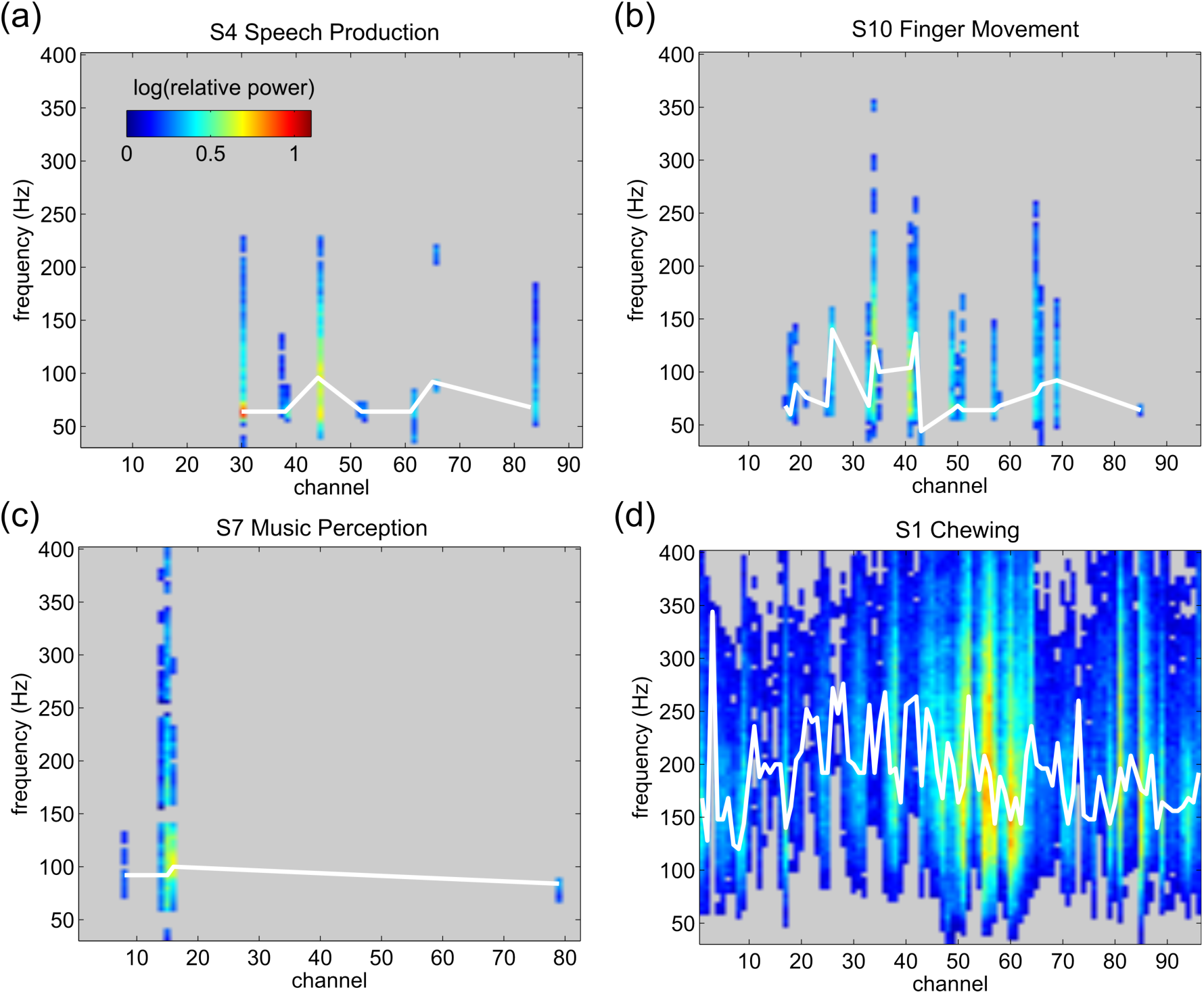
Comparison of task-related (a–c) and chewing-related (d) spectral profiles. The electrodes remained in the original order: first the grid electrodes, then all other intracranial electrodes like strip and depth electrodes. EEG electrodes were excluded from this comparison. The number of electrodes with significant power increases was considerably higher for the ChREs than for the control data. The frequency pattern, quantified by determining the FMP for each electrode (see Table 2 and Figs. 6 & 7), was also different.

**Figure 6:**
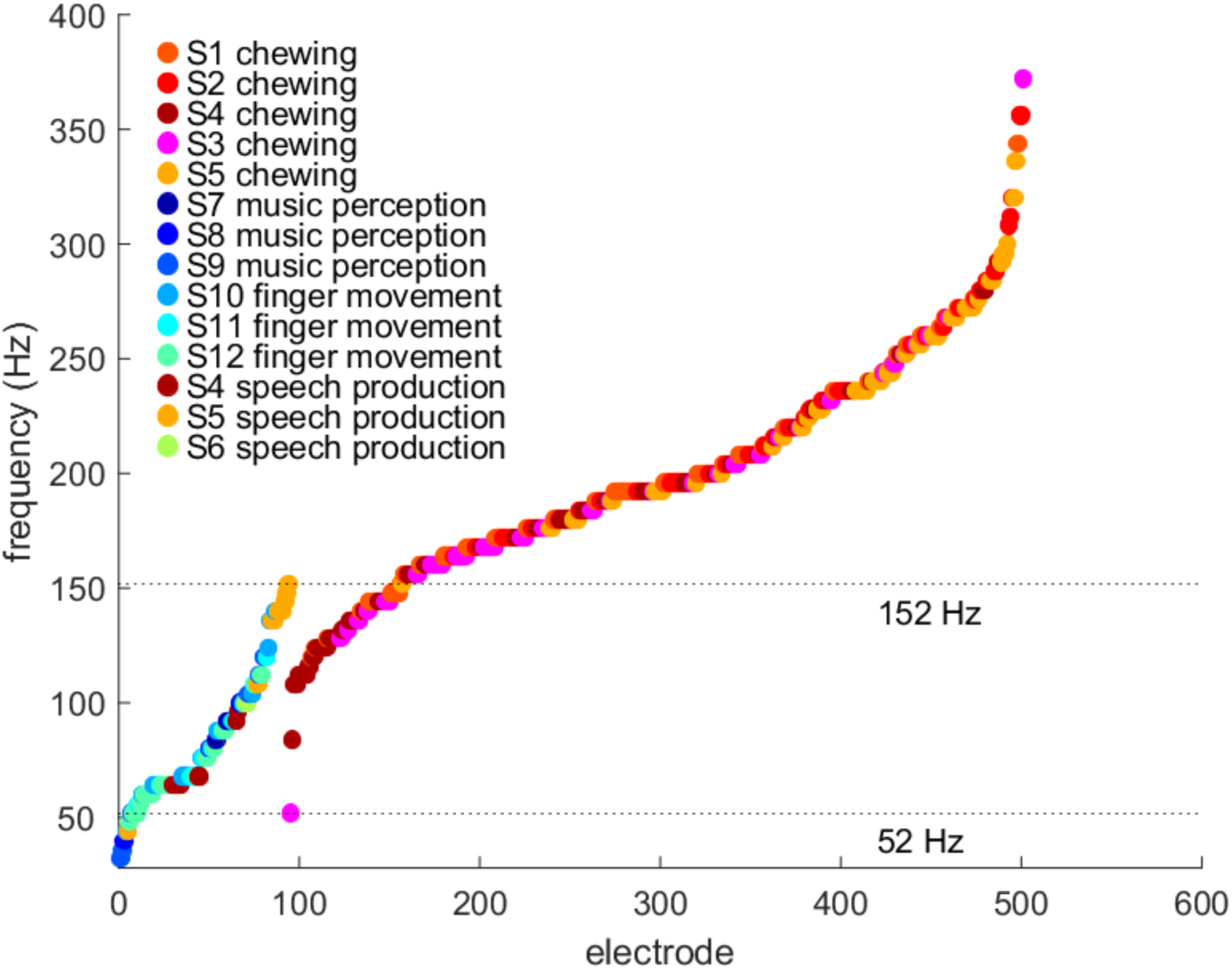
Distribution of FMPs event-related brain activity (left dotplot) and ChREs (right dotplot) across all iEEG electrodesanalyzed in this study. The x axis corresponds to an arbitrary electrode number and the y axis corresponds to the peak frequency (FMP). The values of the FMPs were derived from the results presented in Fig. 5. A unique color was assigned to each subject and the values were sorted by the frequency separately for chewing and control data. Frequencies above 152 Hz were derived uniquely from the ChRE data and frequencies below 52 Hz only from the control conditions.

**Figure 7:**
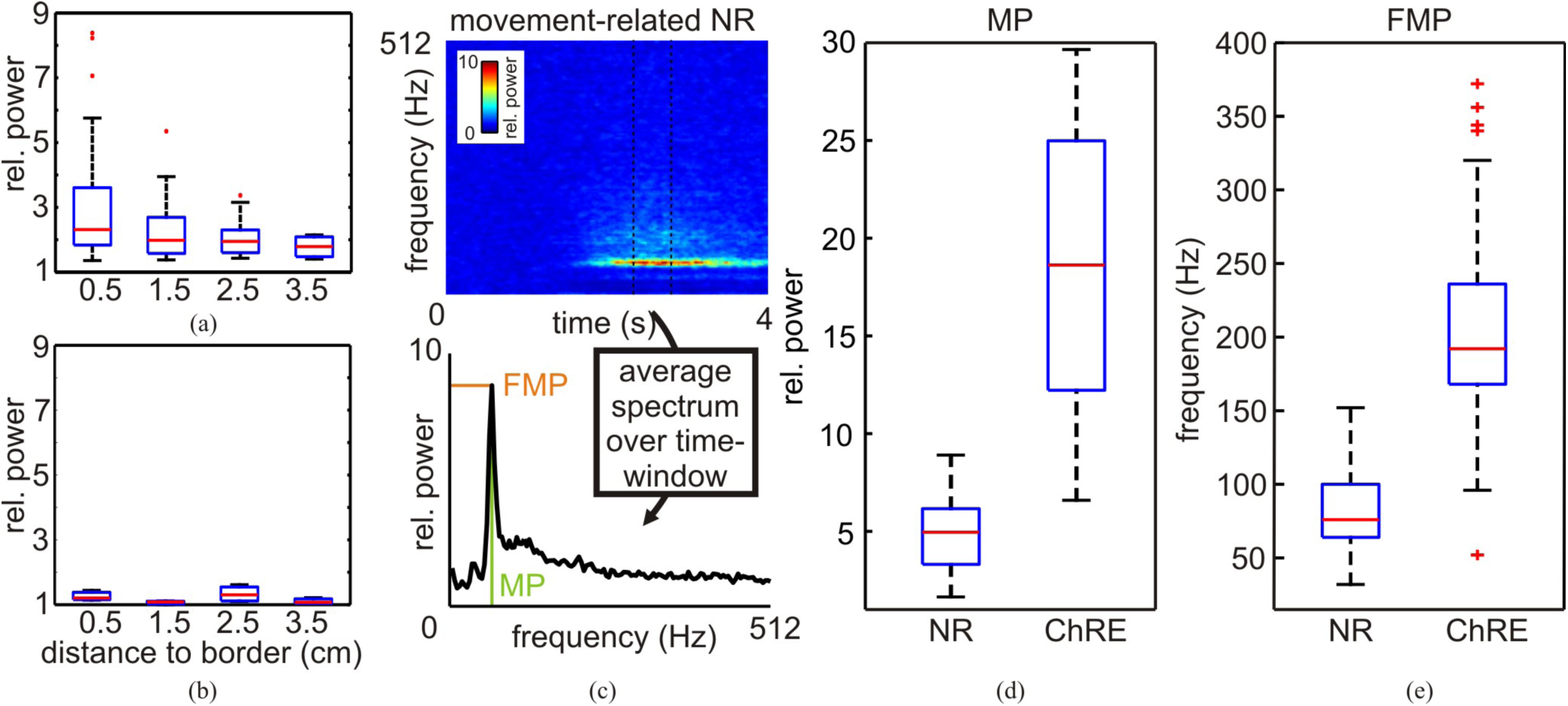
Comparison of topographic and spectral profiles of ChRE and neural iEEG responses. Chewing-related (a), but not neural effects (b) showed the highest relative power at the edge of the electrode grids. **(a**,**b)** Chewing-related relative power decreased as a function of electrode distance to the edges of the grids (S1). This was not the case for ECoG responses of neural origin (b, speech production of S4 as a representative example). **(c)** Illustration of the determination of the maximal power (MP) and the frequency with the maximal power (FMP). A representative neural responses (NR) is shown. **(d**,**e)** Boxplots of the MP and of the FMP, respectively. Red vertical lines indicate the median across subjects (d) and channels (e) (see methods for further details). The box margins indicate the interquartile range (IQR). The whiskers extend to the most extreme value within box margins extended by 1.5 times the IQR, data points outside this range are indicated by red crosses.

**Figure 8:**
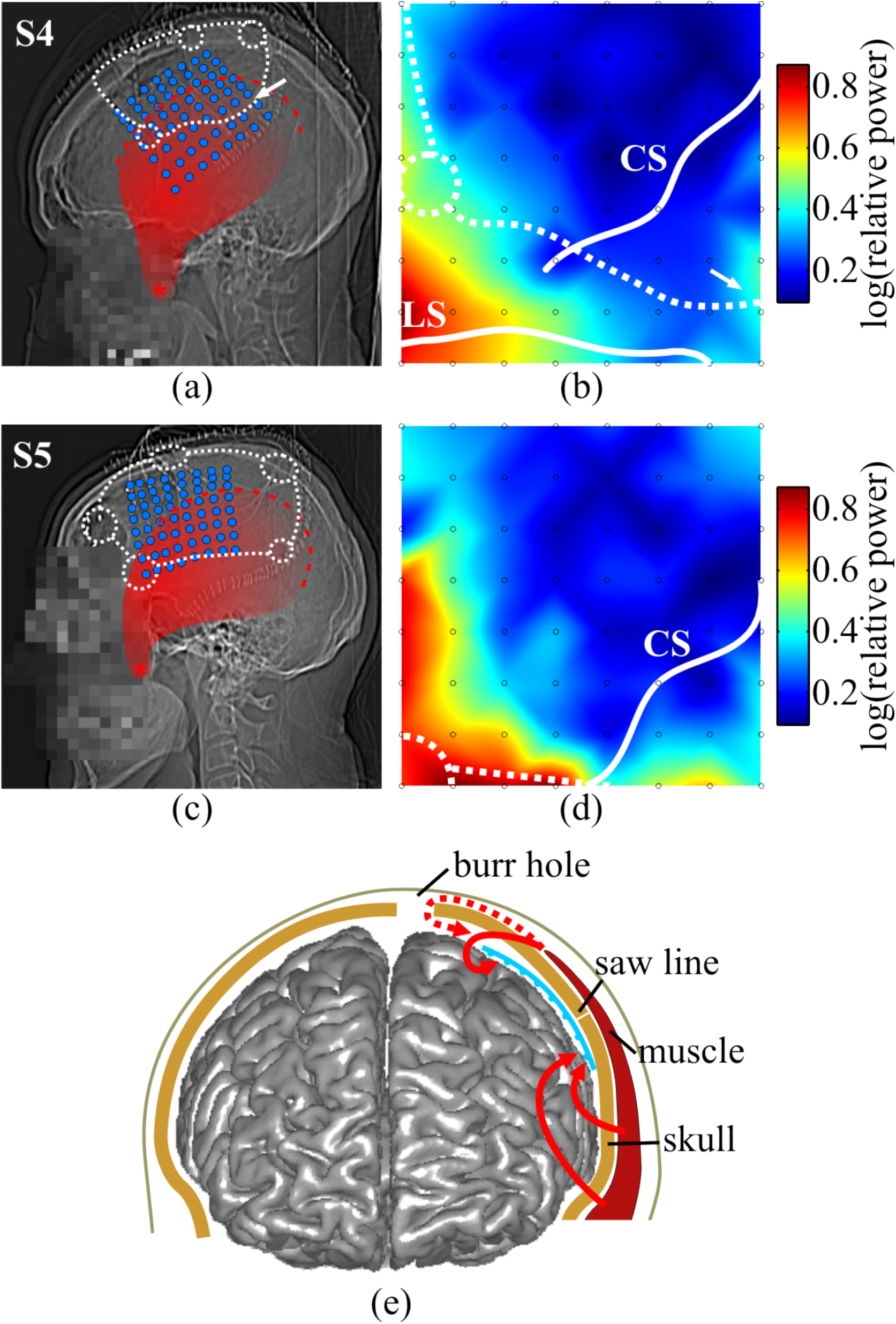
Relation of craniotomy defects to the intracranial ChRE topography. **(a, c)** Lateral X-ray with superimposed positions of implanted electrodes (blue), showing burr holes and saw lines (white dashed lines), and the temporal muscle (red) with the temporal line (red dashed line) as its origin and the coronoid process of the mandibular bone (red asterisk) as its insertion. **(b, d)** The intracranial topography of ChRE in the gamma frequency range (32 to 400 Hz). Electrode positions are marked by gray circles. The saw lines and burr holes are indicated by white dashed lines, the lateral (LS) and central sulci (CS) are indicated by continuous white lines. The white arrow in (b) indicates a power increase on the edge of the grid, possibly caused by the saw line crossing the edge of the grid at this position. **(e)** A summary illustration of extracranial-to-ECoG signal propagation in a schematic anterior view of the brain. The electrode grid (blue) acts as an insulator forcing electric currents to pass around its edges. As the basic spatial pattern of chewing-related signals in the ECoG was largely independent of the individual configuration of craniotomy defects (a)–(d). Thus, the direct pathway through the bone (solid red arrows) plays an important role for extra-to-intracranial volume conduction, in addition to the pathway through the craniotomy defects in the skull (dashed red arrow). (a)–(d) adapted from (Fiederer et al., 2016).

### 2.3 Electrical cortical stimulation mapping

Electrical cortical stimulation mapping (ESM) was performed to identify eloquent cortex using a nerve stimulator (NS 60, Inomed, Emmendingen, Germany) in a constant-current mode. Trains of 7-s duration consisting of 20-ms pulses of alternating-polarity square waves of 200 µs each were applied. Stimulation intensity was increased in steps of 1–2 mA up to 15 mA or until the observation of a sensory, speech-related, or motor effect, whichever occurred first. See also (Glanz (Iljina) et al., 2018; Ruescher et al., 2013) for a more detailed description. The results of electrical stimulation were used to juxtapose the functional topography of body-part related effects with the topography of iEEG responses (Figs. 3 & 4).

### 2.4 Comparison of ChRE with event-related neural activity

For comparison of ChRE (in S1-S5) with iEEG signals of neural origin we referred to the following datasets: in S4, S5 and S6, speaking-related activity during non-experimental, real-life communication was analyzed (Glanz (Iljina) et al., 2018); S7, S8, and S9 participated in a music perception task where the subjects were presented with six-chord sequences (Sammler et al., 2013, 2009); S10, S11 and S12 performed index-finger flexions following (Ball et al., 2004). This sample with different conditions was used to extract general spectral properties of the iEEG responses. For these data, the baseline period was defined as the first 10 time-frequency bins at the beginning of each trial. Because of the different behavioral conditions, a single time window of fixed length cannot be expected to optimally match the timing of all neural responses across all conditions. Therefore, an optimized time window was computed for each subject and condition using the following procedure. First, the median relative power in the gamma band (32–400 Hz) was calculated for all time windows in an interval between 25 and 500 ms and for all possible onsets in the 4-s data epochs. The number of frequency bins with a significant power increase was determined for each condition (false discovery rate (FDR)-corrected (Benjamini and Hochberg, 1995) signtest at a q-level of 0.001 for all tests in the optimization procedure). The parameter combination (window length and onset) which elicited the highest number of significantly increased frequency bins was then selected for further analysis (see also Fig. 2 for a summary on data analysis).

To characterize the spectral profiles of ChRE and of the neuronal signals, we thus calculated the maximal relative power (MP) and frequency of the maximal relative power (FMP) for each electrode (see Figs. 2 & 7c). For the MP, we compared the maximal relative spectral magnitude value per subject (i.e., distribution in Fig. 7d is across subjects. For the FMP, we compared all electrodes in all subjects with significant responses in the ChRE and control conditions (i.e., distribution in Fig. 7e is across electrodes).

To determine the spatial distribution of ChRE in iEEG, relative power changes were averaged over trials, and effects were interpolated across the topography of the electrode grid (Fig. 8). Furthermore, correlation coefficients (Spearman’s ρ) between the relative power and the distance of the electrodes from the border of the grid, for both ChRE and the responses in the control data were calculated (Figs. 7a,b, respectively).

## 3. Results

### 3.1 Topography of intracranial ChREs and event-related neural activity

Topographic results are reported for 8×8 ECoG grids only, because of their large and continuous spatial extent. In agreement with (Fiederer et al., 2016), we show that ChREs are clearly present in the ECoG data and provide additional information about their properties. ChRE were manifested as relative power increases over a broad frequency range at nearly all ECoG electrodes. Effects were stronger at the edges of the grid than in its center, and maximal power increases occurred in the antero-lateral corner of the electrodes grid. This effect is further demonstrated in Fig. 7a, where the distribution of power modulations is shown as a function of electrode distance to the nearest edge of the electrode grid. Correlation coefficients (Spearman’s ρ) between relative power and distance to the edge were S1: r=-0.469; S2: r=-0.580; S3: r=-0. 588; S4: r=-0. 303; S5: r=-0.467. The obtained ρ values are negative as small distances were associated with high power. Note however that this edge-dominated topography was only present in 8×8 ECoG grids and not in smaller ECoG grids. Furthermore, ChRE extended without any interruption over the anatomical borders such as the LS or the CS (Figs. 3 & 4a).

Physiological neuronal activity was, in contrast, considerably more localized. It comprised only a small number of electrodes and showed spatial and functional selectivity. Thus, the finger movement tasks elicited effects in the hand sensorimotor cortex (identified by electrical cortical stimulation), speech production in the articulatory cortex (Fig. 4c), and music perception showed effects in the superior temporal cortex implicated in auditory perception (not visualized).

### 3.2 Spectral characteristics of intracranial ChREs and event-related neural activity

To quantify the influence of chewing-related effects (i.e., mainly EMG artifacts of extracranial origin, see Discussion) on both iEEG and EEG signals, the ratio of the maximal chewing-induced power (MP) increase in iEEG and EEG was determined for all subjects. The power increases were stronger in the EEG that in the iEEG data. They differed on average by a factor of approx. 25 (S1: 28.75; S2: 11.03; S3: 20.12; S4: 16.03; S5:49.70). The MP of ChRE in the iEEG were, on average, larger by a factor of approx. 4 than those related to physiological neuronal activity (Fig. 7d).

To characterize and compare frequency profiles of ChRE with neural iEEG responses, electrodes and frequencies with significant power increases were calculated (Fig. 5). Across all subjects 406/410 electrodes (99.0%, Table 2) of the iEEG electrodes showed at least one frequency bin with a significant power increase related to ChRE (see Table 2). In the frequency range between approximately 150 and 350 Hz, almost all electrodes showed significant power increases related to chewing (Fig. 5j–n). At some electrodes, ChREs spread over the entire analyzed frequency range from 32 to 400 Hz (e.g., Fig. 5d, electrode 48 and 49 in S1). In contrast, only 92 out of 776 electrodes (11.9%) of the electrodes in the control data showed significant power increases (Table 2).

**Table 2:** Properties of the frequencies with the maximal spectral power (FMP) related to chewing (upper part) opposed to the equivalent values related to neuronal activity (lower part). Abbreviations: FMP_max_ maximum of FMP, FMP_med_ its median, and FMP_min_ correspondingly its minimum; Ch: ChRE, Sp: speech production, Mu: music perception, Fi: finger movements; *ch*_*sig*_: number of electrodes with significant power increases; *ch*_*tot*_: total number of iEEG electrodes. The values were derived from the results presented in Fig. 5.

| Subject | FMP<sub>min</sub> | FMP<sub>med</sub> | FMP<sub>max</sub> | ch<sub>tot</sub> | ch<sub>sig</sub> |
| --- | --- | --- | --- | --- | --- |
| S1 Ch | 120 Hz | 192 Hz | 344 Hz | 96 | 96 |
| S2 Ch | 156 Hz | 212 Hz | 356 Hz | 94 | 92 |
| S3 Ch | 52 Hz | 172 Hz | 372 Hz | 64 | 64 |
| S4 Ch | 96 Hz | 168 Hz | 292 Hz | 92 | 92 |
| S5 Ch | 152 Hz | 240 Hz | 340 Hz | 64 | 62 |
| S4 Sp | 64 Hz | 64 Hz | 96 Hz | 92 | 9 |
| S5 Sp | 108 Hz | 140 Hz | 152 Hz | 72 | 10 |
| S6 Sp | 100 Hz | 100 Hz | 100 Hz | 84 | 1 |
| S7 Mu | 84 Hz | 92 Hz | 100 Hz | 82 | 5 |
| S8 Mu | 80 Hz | 92 Hz | 100 Hz | 98 | 2 |
| S9 Mu | 32 Hz | 104 Hz | 120 Hz | 56 | 8 |
| S10 Fi | 44 Hz | 68 Hz | 140 Hz | 96 | 21 |
| S11 Fi | 56 Hz | 76 Hz | 120 Hz | 84 | 12 |
| S12 Fi | 48 Hz | 64 Hz | 112 Hz | 112 | 24 |

To further quantify spectral characteristics the frequency with the maximal power was determined (FMP). Across subjects, the FMP in the iEEG electrodes ranged between 52 and 372 Hz in the ChRE condition and between 32 and 152 Hz in the control data representing neural responses (Figs. 6 & 7e). The FMP derived from chewing-related spectra were generally higher than in the neural responses. Nevertheless, there was an overlap in the frequency range between 52 and 152 Hz (Figs. 6 & 7e). A further distinctive feature between ChRE and neural effects was that broadband gamma increases were only in the latter case typically accompanied by a power decrease in the alpha- and beta-frequency band (8 to 30 Hz (e.g. grid electrode F4 in Fig. 4c).

### 3.3 The intracranial ChRE topography is largely independent of the position of craniotomy defects

The positions of burr holes and saw lines were obtained from CT scans to determine the role of craniotomy-related skull defects (burr holes and saw lines) on the intracranial topography of ChRE. Despite considerably different positions of both burr holes and saw lines, the topography of relative power changes in the gamma frequency range (32 to 400 Hz) remained remarkably constant (Fig. 8b,d). The MP changes in each subject were observed in the antero-lateral corner of the grid, i.e., in a region which was situated directly underneath the belly of the temporal muscle, regardless of whether a burr hole or a saw line was present in this region (Fig. 8a) or not (Fig. 8c). Additionally, strip electrodes located in the direct viscinity of the belly of the temporal muscle had MP changes similar to those of the antero-lateral corner of the grids (e.g. Fig. 5d).

## 4. Discussion

This study is the first comprehensive characterization of non-ocular EMG effects in iEEG based on a larger sample of subjects (in contrast to previous single-case studies, such as (Liu et al., 2004b; Otsubo et al., 2008b), and studies on ocular effects (Ball et al., 2009b; Jerbi et al., 2009b; Kovach et al., 2011b)).

In all subjects, we found ChRE with a broad frequency distribution (from <25 to above 400 Hz, e.g., Fig. 5d) affecting a wide range of cortical areas in the frontal, parietal, and temporal lobes, as well as deeper areas like the inter-hemispheric cleft and the hippocampus.

Further, we contrasted ChRE with neural activity and showed clear distinctions with regard to their spectral and topographic characteristics.

### 4.1 Intracranial ChREs were mainly due to extracranial EMG activity

As we have previously argued (Fiederer et al., 2016), taking the spectral and topographic characteristics into consideration, it appears most plausible to assume that ChREs mainly arise from EMG activity of the masticatory muscles. It is very improbable that ChREs result from neural activity related to the act of chewing. This is supported by two functional magnetic resonance imaging (fMRI) studies that demonstrated focal BOLD signal changes related to chewing, tongue tapping, or swallowing in the primary sensory and motor cortex (Malandraki et al., 2009; Onozuka et al., 2002). This focal BOLD response pattern is in contrast to the ChRE that were spatially widespread, overriding structural and functional boundaries (Fig. 3 & 4a). Furthermore, the frequencies with the highest relative power of the ChRE were, in most cases, higher (25th FMP percentile > 150 Hz) than in any of the investigated examples of control responses representing neural activity (FMP_max_ 152 Hz; Figs. 6 & 7e and Table 2). Then, the spectral profiles of the latter were typically accompanied by low-frequency power decreases, and gamma-band increases (Fig. 4c), a well-established feature of event-related neural-population responses of the cerebral cortex (Crone et al., 1998; Pfurtscheller and Lopes da Silva, 1999), while ChRE showed no low-frequency power decreases (Figs. 3 & 4a). Finally, ChRE power in our EEG data was approx. 25 times higher than in the iEEG, whereas stronger intra-than extracranial effects would be expected for recordings of neural activity. However, as already surmised in our previous study (Fiederer et al., 2016), a weak focal neural signal albeit masked by higher powered extracranial EMG may be present in the ChRE.

### 4.2 ChRE topography is largely independent of craniotomy defects

As expected for signals originating from an extracranial source, the power of the ChRE was much higher (approx. by a factor of 25) in the EEG electrodes than in the iEEG electrodes. Further, the ChRE were spatially widespread, consistent with the spatial low-pass filtering properties of the skull (Nunez and Srinivasan, 2006).

The basic spatial pattern of gamma-band ChRE in our study was observed largely independent of the individual configurations of craniotomy defects (Fig. 8e). We found only limited evidence for a pronounced ‘reverse breach’ effect (personal communication of J. Gotman cited in (Otsubo et al., 2008b)) with focalized responses close to burr holes that might be mistaken for typically focalized physiological, neural event-related effects as shown in Fig. 4c and in previous ECoG studies (Aoki et al., 1999; Crone et al., 1998, 2001a). Nevertheless, increased extra-to-intracranial signal conduction must be expected below craniotomy defects as long as they are not filled by low-conductance material (air, silicone-embedded electrode wires), which is at least as insulating as the surrounding skull. Some of our observed details of the ChRE-related intracranial gamma-band topographies may, indeed, be due to such effects. An example is the increased power at the intersection of the posterior edge of the subdural grid and a saw line in S4 (marked by a white arrow in Fig. 8d). Notably, such cases were restricted to the edge of the grid and did not extend along the whole course of the saw line, moste likely due to the insulating properties of the silicone ECoG grid.

This independence of craniotomy defects in particular, and the intracranial spatial distribution of ChRE in general, may be explained by a model summarized in Fig. 8e: EMG activity from masticatory muscles passes through both craniotomy defects and intact parts of the skull and around the insulating substrate of the ECoG grid, which leads to stronger effects near the edges of the grid. The overall spatial distribution is smooth due to the extended size of the generating muscles and the dominating effect of currents passing through the skull with spatial low-pass filtering properties (Nunez and Srinivasan, 2006). The strongest ChRE in the antero-lateral corner of the ECoG grids in all investigated subjects can be explained by its proximity to the belly of the temporal muscle (Fig. 8a–d), one of the main active muscles during chewing. This spatial relationship was also confirmed in 3 subjects having strip electrodes located in the direct viscinity of the belly of the temporal muscle. In the opposite direction (intra-to-extracranial, i.e. from neural activity to EEG), previous simulations indicated that substantial spatial distortions of scalp surface EEG are to be expected due to the fact that currents have to pass around the insulating material of the silicone ECoG grid (Lanfer et al., 2012a; Nunez and Srinivasan, 2006; von Ellenrieder et al., 2014; Zhang et al., 2006). These assumptions were also supported by the results of our previous study on volume conduction simulations in a single subject (Fiederer et al., 2016), where we showed that the ChREs indeed mainly propagate directly through the skull. Furthermore, the simulations in the aforementioned study also confirmed that the silicone ECoG grids substantially attenuated and distorted the measured ChREs. Without craniotomy defects and without grid, simulated ChREs were 21% stronger than in our measurements. By contrast, removing craniotomy defects and keeping the silicone grid decreased ChREs by only 6%. The silicone grid thus attenuated ChREs by 27%.

In the data related to neuronal activity, only a small number of electrodes showed clear task-related neural responses (approx. 12 % of all iEEG electrodes, Table 2), and the effects were typically confined to small, focalized areas (Figs. 3 & 4c). For the speech production task, it was further shown, that the relevant electrodes were located in the areas that elicited orofacial responses during electrical cortical stimulation (Fig. 4b,c). Moreover, in contrast to the ChRE, the effects related to neuronal acitivity were not concentrated on the border of the electrode grids, as we are demonstrating by correlating the effect with the distance to the border of the grid (Fig. 7a vs. b).

### 4.3 ChREs involve a broader frequency range than neuronal activity

In previous reports on surface EMG of facial and masticatory muscles, EMG was characterized by signal increases in a broad frequency range from several Hz to over 500 Hz, with peak frequencies between 80 and 160 Hz for the temporal muscle ((Boxtel et al., 1983): peak frequency ∼130 Hz; (Palla and Ash Jr, 1981): ∼80 Hz; (Yuen et al., 1989): ∼160 Hz), matching well the ChRE spectra we observed extracranially in the present data (Fig. 1b,c). As extra-to-intracranial volume conduction is largely frequency-independent (Nunez and Srinivasan, 2006), we expected a similar spectral profile in intracranial data. Consistent with this expectation, ChRE in iEEG involved a broad frequency range from <25 Hz to above 400 Hz (e.g., Fig. 5d, limited by the anti-aliasing filters and sampling rate of our recordings). Across all electrodes, the frequencies with the maximal iEEG ChRE power increases were observed in a range of approx. 100–350 Hz (Fig. 6 & 7e). However, in a study which analyzed the masseter and temporalis muscles, as well as five other facial muscles in terms of the spectral properties of their activity, the temporal-muscle EMG spectrum was found to be atypical in comparison with the other muscles in the sense that it had an exceptionally high peak frequency, whereas the peak frequencies of other facial muscles (e.g., buccinator or frontalis) were below 50 Hz (Boxtel et al., 1983). Thus, artifacts related to facial expressions or speaking, and hence originating predominantly from mimic muscles may be also present in frequency bands other than those characterized in this study.

In the event-related datasets, we found a differential spectral pattern. In the whole frequency spectrum, a pattern of power increases in the gamma band, accompanied with a decrease in lower frequencies, was found. The MP increases related to chewing were higher by a factor of approx. 4 than those related to neuronal activity. While ChREs affected virtually the entire frequency range nearly all of the electrodes, power modulations related to the control data rarely exceeded frequencies of 250 Hz (only 12 out of 92 electrodes showed significant effects in frequencies above 250 Hz in all nine subjects performing experimental tasks, see Fig. 5c,d for 2 examples). Further, the FMP was lower for the event-related data than for the ChRE, albeit with an overlap at some electrodes in the range between 52 and 152 Hz. Of note, the highest values for the FMP were obtained from one subject in the speech production condition (Fig. 6). We argue that these still reflect neural activity as the response was spatially focal (10 contacts), located at the center of the grid (cf. Table 2 and Fig. 6), and largely co-localized with the ESM-defined mouth sensorimotor cortex. Still, although we assumed that the effects in the task-related datasets to be predominantly of neural origin, it can not be excluded that a certain amount of EMG artefacts may be present here as well. Taking together the topographic and spectral characteristics, a more finegrained nature of the neuronal activity as compared to the ChRE became apparent.

### 4.4 Increasing the spatial extent of the silicone edges may improve the quality of ECoG recordings

We have shown that both ECoG data (this study) and simulations (Fiederer et al., 2016) support the concept that the silicone sheet, in which ECoG electrodes are embedded, attenuates the signal of extracranial electric sources. Thus, the silicone sheet actively contributes to the signal quality of ECoG. However, this shielding effect scales with the distance of the electrodes to the edges of the grid and is thus neither present in smaller grids nor in strips. Extending the edge of the silicone sheet may thus further increase the shielding effect and consequently further increase the robustness of ECoG recordings against, e.g., EMG artifacts.

However, there are indications that the ECoG grid design may have an effect on the occurrence of clinical complications (Hamer et al., 2002; Önal et al., 2003; Wiggins et al., 1999; Wong et al., 2009). According to these studies, the following parameters were repeatedly found to increase the occurrence of clinical complications significantly: larger ECoG grids, increased total number of electrodes, longer implantation durations and left hemisphere implantation. When looking closely at the definition of ECoG grid size, it becomes apparent that it refers to the number of electrodes contained in an ECoG grid (Önal et al., 2003; Wong et al., 2009), and not its area. Furthermore, Wong and colleagues argue that the increased stiffness and thus increased pressure on vessels is the main factor behind the increased occurrence of clinical complications when implanting ‘larger’ ECoG grids (Wong et al., 2009). It is thus still an open question whether broader borders of an ECoG grid would increase the occurrence of clinical complications or not. Given that one important property of silicone is its great mechanic flexibility, increasing the width of the ECoG grid’s silicone edges without additional metal components may reduce its overall stiffness. Thus, enlarging the silicone border of an ECoG grid may increase the signal quality without compromising patient safety.

## 5. Conclusion

By comparing intracranially-recorded chewing-related events (ChREs) with physiological neural activity underlying several tasks in epilepsy patients, we were able to demonstrate their distinct topographic and spectral characteristics. The differences in topography and in preferred frequencies that we found may be useful for further development of methods for the reduction of EMG artifacts in iEEG. For instance, spatial high-pass filtering may be beneficial to differentiate focalized neural activity from widespread EMG. The silicone grids used in ECoG measurements produced a remarkable shielding effect, which may have implications for the design of future ECoG electrodes and implanted BMI recording devices.

## Acknowledgment

Funding: This work was supported by the German Federal Ministry of Education and Research (BMBF) grants 01GQ0420 to BCCN Freiburg, 0313891 GoBio, 16SV5834 NASS, 01GQ1510 OptiStim, and German Research Foundation (DFG) grant EXC 1086 BrainLinks-BrainTools.

